# Recent mixing of *Vibrio parahaemolyticus* populations

**DOI:** 10.1101/354761

**Authors:** Chao Yang, Xiaoyan Pei, Yarong Wu, Lin Yan, Yanfeng Yan, Yuqin Song, Nicola Coyle, Jaime Martinez-Urtaza, Christopher Quince, Qinghua Hu, Min Jiang, Edward Feil, Dajin Yang, Yajun Song, Dongsheng Zhou, Ruifu Yang, Daniel Falush, Yujun Cui

## Abstract

**Background:** Humans have profoundly affected the ocean environment but little is known about anthropogenic effects on the distribution of microbes. *Vibrio parahaemolyticus* is found in warm coastal waters and causes gastroenteritis in humans and economically significant disease in shrimps.

**Results:** Based on data from 1,103 genomes, we show that *V. parahaemolyticus* is divided into four diverse populations, VppUS1, VppUS2, VppX and VppAsia. The first two are largely restricted to the US and Northern Europe, while the others are found worldwide, with VppAsia making up the great majority of isolates in the seas around Asia. Patterns of diversity within and between the populations are consistent with them having arisen by progressive divergence via genetic drift during geographical isolation. However, we find that there is substantial overlap in their current distribution. These observations can be reconciled without requiring genetic barriers to exchange between populations if dispersal between oceans has increased dramatically in the recent past. We found that VppAsia isolates from the US have an average of 1.01% more shared ancestry with VppUS1 and VppUS2 isolates than VppAsia isolates from Asia itself. Based on time calibrated trees of divergence within epidemic lineages, we estimate that recombination affects about 0.017% of the genome per year, implying that the genetic mixture has taken place within the last few decades.

**Conclusions:** These results suggest that human activity, such as shipping and aquatic products trade, are responsible for the change of distribution pattern of this marine species.

## Background

Hospitable environments for particular marine microbes can be separated by large distances but whether dispersal barriers substantially influence their distribution and evolution is unknown. There are many studies of distribution of marine microbes e.g. [1-4], but these typically survey patterns of macro-scale diversity. Differences in species level or genus level composition between locations are as likely to reflect environmental heterogeneity as dispersal, making the patterns difficult to interpret. Recent spread of microbes between continents has been documented for lineages that cause pathogenic infection of humans, including notorious clonal groups within *Vibrio parahaemolyticus* and *Vibrio cholerae* [5-8]. However, these lineages are unusual in using humans as vectors, which might facilitate long-range dispersal as in the case of the Haitian cholera outbreak [9]. We currently have little information on rates of spread of the great majority of environmental organisms that do not colonize large-animal hosts.

*V. parahaemolyticus* prefers warm coastal waters and causes gastroenteritis in humans [10, 11]. Disease outbreaks became common from 1990s and became global, due to spread of particular clones which are responsible for the great majority of recognized human infections [5], which has been attributed to factors such as El Niño and climate change [12-14]. It is not clear to what extent this pattern is historically typical, or whether it instead reflects better surveillance and different patterns of usage of marine resources. These clones also make up a small fraction of the *V. parahaemolyticus* diversity and only a very small fraction of strains isolated during environmental sampling.

The *V. parahaemolyticus* genome undergoes high rates of homologous recombination with other members of the species [15, 16]. We have previously found evidence that the species is split into several populations [16]. Members of a population are not necessarily particularly related at the clonal level, for example they may have recombined their entire genomes since sharing a common cellular ancestor, but they are nevertheless on average more similar to each other than to members of other populations because they have acquired DNA from a common gene pool. Previously we found evidence of a single population with a well-mixed gene pool in Asian waters and for one or more differentiated populations in the US [16].

Here we use a larger and more broadly sampled collection of 1,103 genomes to examine the global population structure of the species. We find four populations with different but overlapping modern geographic distributions as well as a small number of hybrid strains. Under the assumption that genetic exchange between strains is constrained by geography, the current extent of overlap is too high to maintain the populations as distinct entities and we conclude that most of this mixing is likely to have taken place within the last few decades, possibly coinciding with the recent emergence of pandemic clones.

## Results and Discussion

### Distribution of *V. parahaemolyticus* populations

We analyzed genomes of 1,103 strains including 392 new strains sequenced as part of this study. These strains were isolated from a mixture of sources during 1951-2016, and covered 24 countries (Supplementary Fig. 1 and Supplementary Table 1). Clonal relationships between strains can be inferred from identifying long stretches of near-identity, corresponding to regions of the genome that have been inherited by direct descent since the strains shared a common ancestor, or, more simply, by the strains having a small number of SNP differences between them genome wide, which can be revealed by the Neighbor-Joining (NJ) tree (Fig. 1a). Based on criterion of high nucleotide identity, the dataset contains 13 clonal groups, with 10 associated with human disease and 3 associated with the environment (Supplementary Table 1).

**Figure 1.**
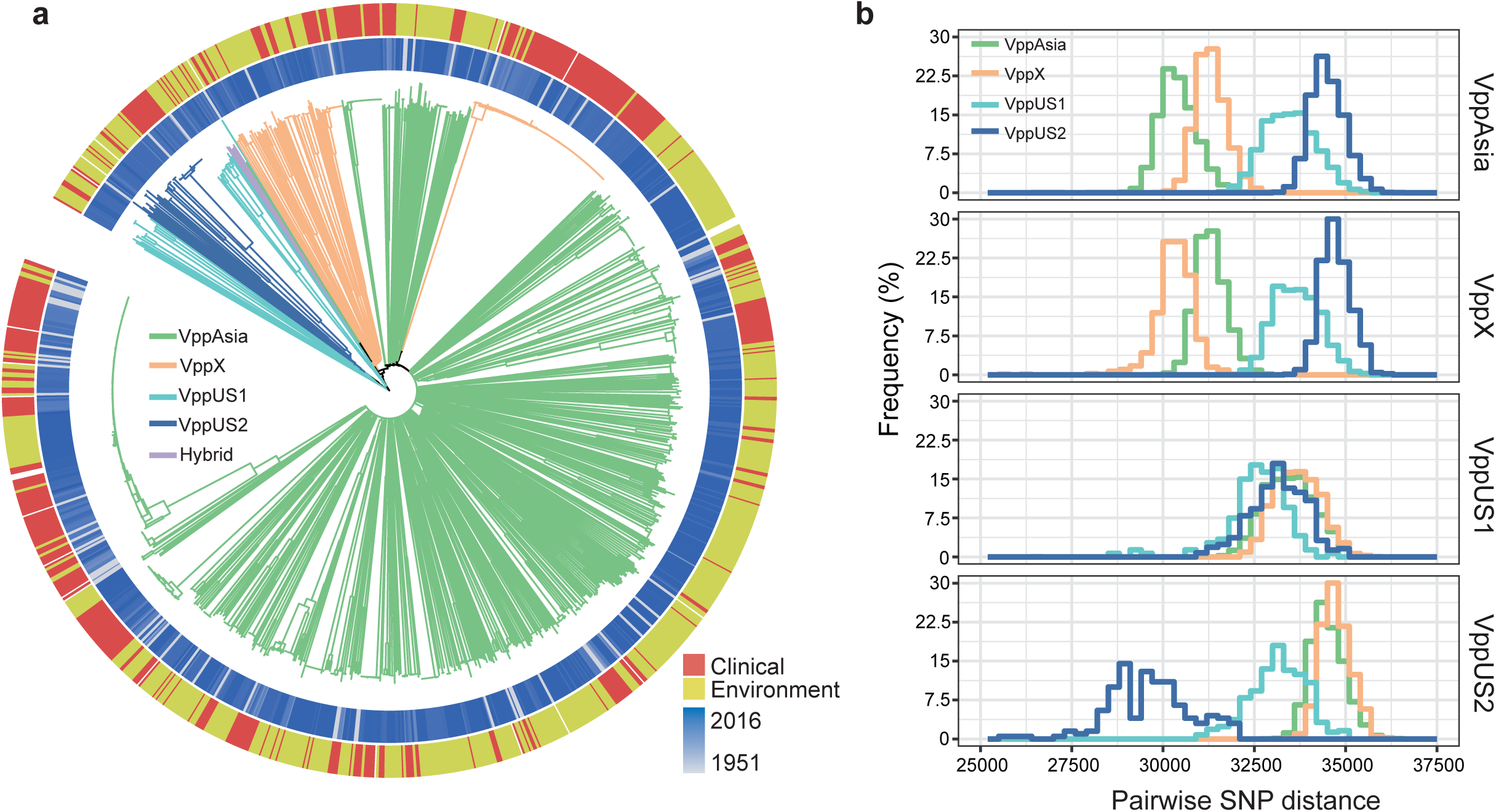
Population structure of *V. parahaemolyticus* and relationships within and between populations. (a) NJ tree of 1,103 *V. parahaemolyticus* stains based on 462,214 SNPs. Branch colors indicate populations defined by fineSTRUCTURE, green for VppAsia, orange for VppX, light blue for VppUS1, dark blue for VppUS2, purple for hybrid strains. The ring colors from inner to outer indicate isolation time and sample type, respectively. The blank indicates information not available. (b) SNP distance within and between populations based on 469 non-redundancy strains. Colors indicate populations and are consistent with branch colors of panel a.

The presence of clonally related strains in the data complicates analysis of deeper population structure, so we first removed closely related isolates to make a “non-redundancy” dataset of 469 strains, in which no sequence differed by less than 2,000 SNPs in the core genome (Methods). We used fineSTRUCTURE to identify distinct populations [17]. In total, 115 populations were identified in this initial analysis, however most comprised only two or three strains (Supplementary Fig. 2a). These are likely to be sets of strains that are clonally related, so we removed all but one from each group and reran fineSTRUCTURE. After several iterations of the same procedure, we identified four populations with between 10 and 217 members and two singletons (Supplementary Fig. 2b). These singletons might be hybrids or representatives of otherwise unsampled populations.

The current distribution of the populations is shown in Fig. 2a. The great majority of isolates from Asia (574/600) are assigned to VppAsia, with all but one of the remainder (VppUS1, isolated from a shrimp farm in Thailand) being assigned to VppX. VppUS1 is found almost entirely in the US and is most common in the Mexican Gulf, with 13 out of the 29 VppUS1 strains are isolated from there. VppUS2 is most common on the US Atlantic coast (20 of 42) and has also been isolated several times in Northern Europe. VppX is most common on the Pacific coast and the Northern part of the US coast. These patterns are not predominantly determined by the spread of human disease clones, since similar patterns are observed if the dataset is restricted to the 469 non-redundancy strains (Supplementary Fig. 3a). The distribution of CG1, the pandemic clonal group that mostly belongs to sequence type (ST) 3 [5], is similar to that of other VppAsia isolates (Supplementary Fig. 3b), while CG2 (ST36), an epidemic group that is abundant in US and Canada [8], has a similar distribution to that of VppX isolates, except that it has not been isolated from Asia (Supplementary Fig. 3b). The distribution of the four populations is also similar when analysis is restricted to strains from the human disease (Supplementary Fig. 3c) or environment (Supplementary Fig. 3d).

**Figure 2.**
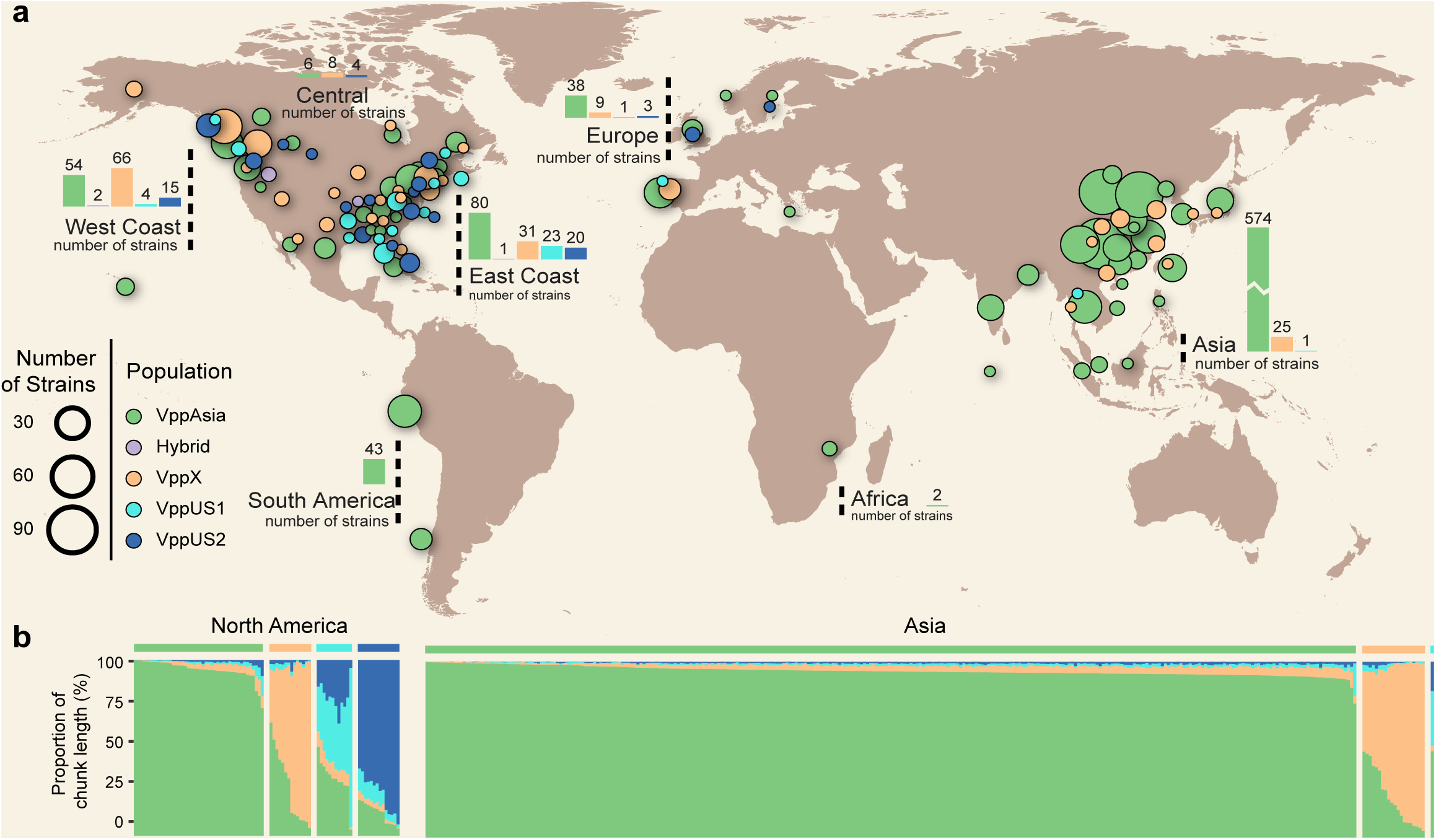
Geographical distribution and admixture of *V. parahaemolyticus* populations. Colors in circle and bar plot indicate populations and are as in Figure 1. Each circle indicates the population composition of a city/country, with radius in proportion to the sample size. Bar plot indicates the ancestry composition inferred by chromosome painting of two geographical regions: Asia and North America. Each vertical bar represents one non-redundancy strain and the proportion of color indicates the contribution of each population. Different populations are separated by blank vertical bar. Only strains with information of isolation location are included in panel a (n = 1,008) and panel b (n = 422).

The 1,103 genomes in this study have been collected for a variety of different purposes and do not represent a defined environmental or epidemiological cohort. Furthermore, sampling numbers in most locations are small and the coasts of Africa and Australia, for example are almost entirely unsampled. Nevertheless, our results demonstrate that at a global scale, geographic distributions of populations overlap considerably and that there is a substantial difference in the frequencies of the populations in the waters of Asia and those of the US Coast (Fig. 2a).

### Relationships amongst populations

The populations have a modest level of differentiation at the nucleotide level (Supplementary Table 2), with *F*_*st*_ values of around 0.1 approximately equivalent to that between humans living on different continents [18], implying that most common polymorphisms are shared between populations. VppUS1 is the most diverse and isolates are no more similar to each other in terms of mean SNP distance than they are to members of the other populations (Fig. 1b). However, according chromosome painting, which is based on haplotype similarity and therefore more sensitive in detecting sharing of DNA due to common descent, all of the members show substantially higher coancestry with other members of the population than any of the other isolates in the dataset (Fig. 2b), implying that the population consists of isolates that share ancestry, rather than being a collection of unassignable genomes. The other populations have consistently lower distances with members of their own populations and VppX and VppAsia are more closely related to each other than they are to VppUS1 and VppUS2.

One explanation for the high diversity of VppUS1 is that it has frequently absorbed genetic material from other populations. In order to test this hypothesis, while avoiding the effect of clonal relationships within the population itself on estimates of relationships with other populations, we painted the chromosomes of each of its members, using the members of the other three populations as donors. A high diversity of painting palettes was observed from VppUS1, with between 43% and 74% assigned to VppAsia and between 15% and 49% to VppUS2 (Supplementary Fig. 4a). By contrast, the other three populations showed lower levels of variation in assignment fractions in analogous paintings (Supplementary Fig. 4b-d). Thus, VppUS1 owes its high diversity to being a hub for admixture, with input from both VppUS2 and VppAsia. The members of VppUS1 in our sample are all clearly distinct in ancestry profile from members of other populations (Fig. 2b), justifying the distinct population label, but if gene flow levels were higher, it seems likely that the population would lose its distinct identity and ancestry patterns would be better described by a continuum than discrete population labels.

### Recent mixing of *V. parahaemolyticus* populations

The observation of distinct populations is informative about patterns of migration in the past. Population genetic theory implies that differentiation between demes can only arise and persist if levels of migration between them are low, specifically on the order of magnitude of one migrant per generation or less [19]. The intuition behind the theory is that once a migrant arrives in a deme, it progressively imports DNA from other strains and becomes more and more similar to the other strains in its new deme. If too many strains are migrants, the demes will progressively lose their distinct genetic profiles and merge into a single gene pool. This theory has been developed for outbreeding eukaryotes [20] and bacterial populations deviate from several of the assumptions of the theory, in ways that are currently not well understood, making quantitative predictions impossible. Nevertheless, the qualitative expectation is that at equilibrium most isolates should have the ancestry profile of the region, with only a small fraction of the isolates having part ancestry from other locations.

The data differs from the qualitative predictions of migration-drift equilibrium because while there are few strains of clearly intermediate ancestry in the dataset, many locations have multiple strains from two or more of the four distinct populations that we have identified, making it not obvious what deme they belong to. Asia is clearly the most likely ancestral home range of VppAsia based on its high prevalence there but it is difficult to define boundaries of likely ancestral ranges for the other three populations with any confidence because the isolates assigned to those populations are too dispersed and they do not make up a clear majority anywhere. Thus the current distribution is qualitatively inconsistent with migration-drift equilibrium.

There are a number of factors which can in principle maintain subdivision when members of more than one populations are found in the same location over long time periods. For example, it is possible that the mechanism by which recombination occurs results in import occurring preferentially from members of the same population. For example, barriers to recombination due to homology dependent mismatch repair has been proposed to account for the differentiation between phylogroups of *Escherichia coli*, despite high overall level of recombination [21] because the mechanism preferentially aborts recombination events between members of different phylogroup. Other mechanisms that can generate barriers to gene flow are strain specific phage, or differences in an ecological niche. However, the pattern of sharing of diversity is very different in *V. parahaemolyticus* to that found in *E. coli*, with high nucleotide diversity and low differentiation between them and there are few highly differentiated loci anywhere within the core genome (Supplementary Fig. 5). It is difficult to conceive of a mechanistic barrier encoded within the genomes or their phage that would effectively constrain recombination between populations enough to explain the low number of hybrids within the dataset as a whole, while also allowing the frequent recombination required to create the freely mixed gene pools we see within populations.

We propose instead that the population is far from equilibrium because barriers to movement of strains have reduced recently. Under this hypothesis, it should be possible to approximately estimate the timescale on which mixing has taken place, based on the amount of introgression found in locations where the different populations now co-occur. Specifically, within our dataset, it is natural to compare the VppAsia isolates within Asia and in North America. Since Asia has been least affected by between continent migration (Fig. 2a), we predict that the VppAsia isolates in North America should have more ancestry from other sources, that they have acquired recently in their new locations. This prediction is borne out, a number of North America VppAsia isolates have high levels of VppUS1 and VppUS2 ancestry and on average the North America VppAsia isolates in the non-redundancy set of 469 strains have 1.01% more (in average 2.97% in North America vs 1.96% in Asia) of their painting palette from VppUS sources than those from Asia (Fig. 2b and Fig. 3a).

**Figure 3.**
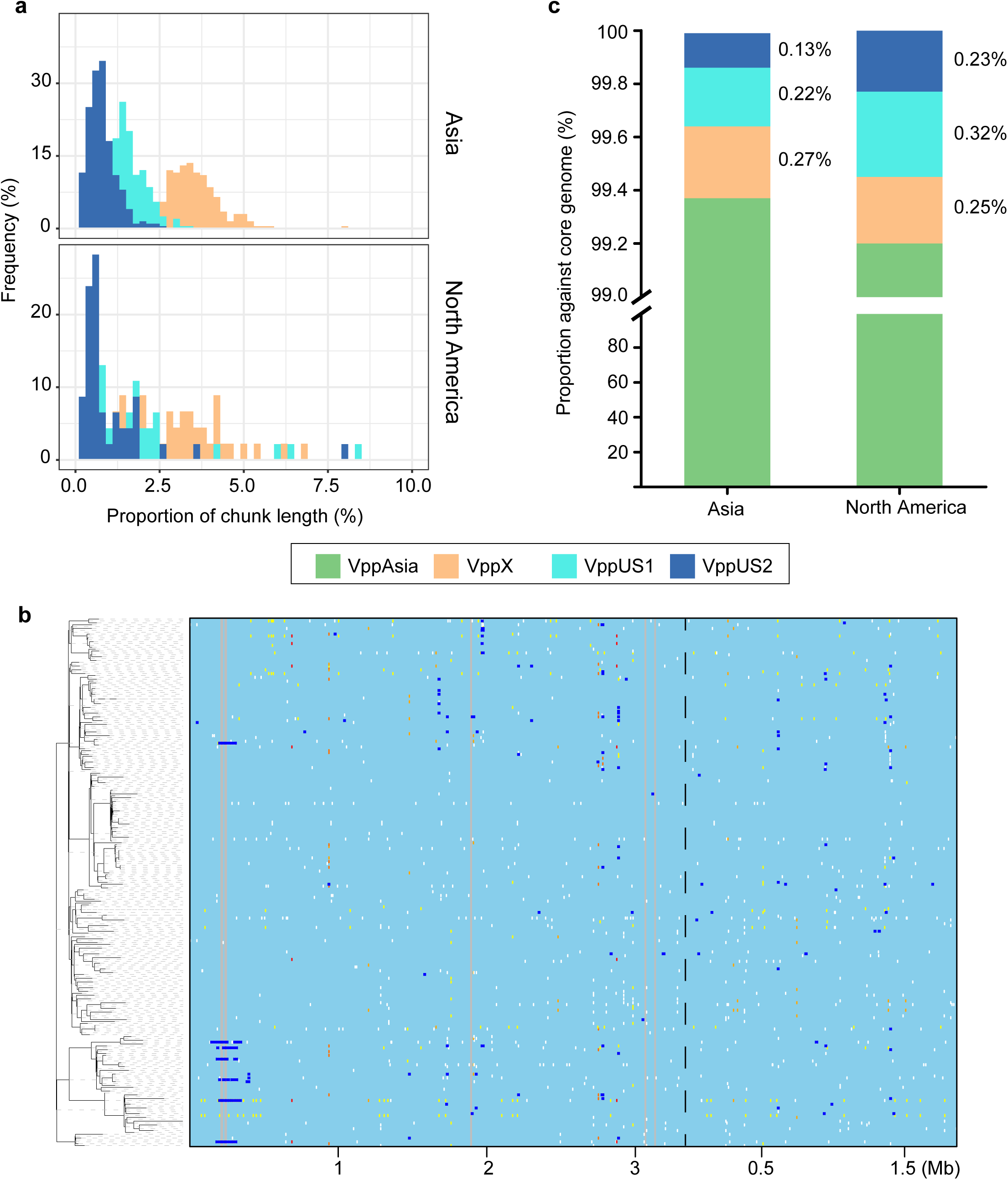
Recent mixing of *V. parahaemolyticus* populations. (a) Ancestry composition of three other *V. parahaemolyticus* populations in VppAsia strains in different geographical regions. The contribution from other populations to the VppAsia is inferred by chromosome painting. X axis indicates the proportion of contributed chunk length of a population in one strain and Y axis indicates the corresponding frequency. (b) ClonalFrameML recombination analysis of 141 CG1 strains. Left: ClonalFrameML reconstructed phylogeny. Right: dark blue horizontal bars indicate recombination events, grey areas indicate non-core regions. Two chromosomes are separated by dot line. (c) Source of recombination fragments of CG1 strains in different geographical regions. Y axis indicates the proportion of recombination fragments input from different population against core genome. Colors in (a) and (c) indicate four populations and are as in Figure 1.

In order to provide a timescale for the acquisition of non-Asian ancestry, we examined the evolution within the largest two clonal populations, CG1 and CG2. We removed recombination regions, then ran BEAST [22] to estimate a clock rate of 5.5 × 10^-7^ per site per year, with very similar values for the two clonal complexes (Supplementary Fig. 6). There are about 313 bases exchanged per mutation (Supplementary Fig. 7), so this implies a rate of recombination of 1.7 × 10^-4^ per site per year. Thus if all of the import into the VppAsia bacteria was from US populations, then it would imply it would take about 59 years (with extreme lower and upper boundaries of 32-151 years) to acquire an extra 1.01% ancestry at this rate of import.

We also examined the origin of imports within CG1, the global pandemic clonal group. As for the VppAsia isolates, a higher fraction of the imports was from the two US populations amongst the isolates found in the North America than for the isolates found in Asia itself. This small difference in ancestry, corresponding to about 0.19% of the genome in total (Fig. 3c), has arisen during around 20 years since the beginning of global spread of CG1 in 1996.

These observations are consistent with a hypothesis that barriers to migration have become substantially weaker within the last few decades, but do not constitute direct evidence that patterns of gene flow between populations have changed. This hypothesis is empirically testable although we do not have a suitable strain collection to facilitate it. For example, if the Asian bacteria have arrived in large numbers in the US recently, then DNA from VppAsia bacteria should make up a higher proportion of recent genetic imports than older ones.

In order to explain why the pattern of dispersal has changed recently, it is necessary to first postulate reasons why dispersal was previously limited. We hypothesis that spread of bacteria between oceans is limited by large distances between environments that are hospitable, making it rare that bacteria survive transportation between them. Large mammals, seabirds and other aquatic organisms travel large distances but do not necessarily provide habitats that *V. parahaemolyticus* can colonize for the days or weeks required to get from one ocean to another. Thus, we propose that dispersal between oceans did occur but was rare.

Humans have changed several aspects of the ocean environment, creating new habitats through effluent discharge, warming and acidifying the oceans through climate change, providing new mobile habitats on the hulls of ship and in ballast water and transporting copeopods and other marine organisms deliberately to facilitate aquaculture or more accidentally through trade in marine products [23, 24]. Several of these could have facilitated transmission of bacteria between oceans. Furthermore, *V. parahaemolyticus* can adapt to colonize copeopods [25] so that for example human-associated dispersal of species such as the manila clam from Asia to the America and Europe [26] could be responsible for the high frequency of Asian *V. parahaemolyticus* there. A single introduction via ballast water or introduction of shellfish for aquaculture would typically have low values of propagule pressure (a single event with few individuals), while recurring introductions through recently increased human activity may contribute in a regular basis introducing trans-ocean migration of *V. parahaemolyticus*.

Further work is required to narrow down the most important factors. To identify the frequency of *V. parahaemolyticus* reads in extensive metagenomic sampling of the open ocean would provide knowledge on natural transmission of this bacterium. One objection to a direct human dispersal, rather than for example a role for climate change is that the absolute number of bacteria transported by ships or trade is likely to be small. However, this objection does not seem especially compelling. The absolute number of bacteria transported from one ocean to another does not need to be very large; if bacteria are fit in their new environment, they can multiply rapidly to constitute a substantial proportion of the bacteria in their new habitat.

## Conclusions

Our results support our earlier conclusion that *V. parahaemolyticus* is subdivided into distinct geographical populations. We have identified 4 clearly differentiated populations, two of which appear to have foci in the US (VppUS1 and VppUS2). A third is predominant in Asia, while the ancestral home range of the forth VppX is difficult to guess based on current sampling. However, these ranges pose a puzzle, in that they overlap substantially, both for environmental and human disease causing isolates, which show approximately similar patterns of distribution. Hybrids are rare, for example, amongst VppAsia isolates found in the US, most have ancestry profiles indistinguishable from strains found in Asia, while a handful have less than 10% introgression from either of the two US populations. The simplest and most parsimonious explanation is that previous barriers to migration have been reduced recently, allowing bacteria to disperse rapidly between continents but that because bacterial recombine relatively slowly (about 0.017% of their genome a year on average), there has not had sufficient time to generate hybrids.

These results have two major implications. Firstly, they suggest that recent human activity has disrupted long-standing barriers to genetic exchange in the oceans and that this has affected microbial population structure. Secondly, changing global patterns of *V. parahaemolyticus* disease incidence may be directly connected to changes in dispersal of the species, rather than being specific to the small number of clonal lineages that are responsible for most of the major outbreaks.

## Materials and Methods

### Bacterial strains

Totally 1,103 strains were used in this research, including 392 newly sequenced and 711 publicly available strains (Supplementary Table 1). The newly sequenced strains were isolated in China during daily food surveillance in 2014. The remaining 711 publicly available strains were downloaded from the NCBI database. The genomes of newly sequenced strains are available in GenBank with the accession numbers listed in Supplementary Table 1.

New sequenced strains were cultured in the LB-2% NaCl agar at 37 °C, and classical phenol/chloroform method was used to the extract genomic DNA.

### Sequencing and assembly

The whole genome DNA was sequenced by using Illumina Hiseq 4000. The pair-end sequencing library with average insert size of 350 bp were build according to the manufacture’s introduction (Illumina Inc., USA). The read length is 150 bp and in average 500 Mb raw data were generated for each strain, which is corresponding to the sequencing depth of approximately 100 fold. The adaptor sequence and low quality reads were filtered and the clean reads were assembled by using SOAPdenovo v2.04 [27] as described before [16]. The number of contigs and average size of assemblies are 263 and 5.1 Mb, respectively.

### Variation Detection

The SNPs were identified by aligning the *V. parahaemolyticus* genomes against with the reference genome (RIMD 2210633) by using MUMmer [28] as previously described [16], and only bi-allelic SNPs were used in further analysis. As the number of detected SNPs would relate with core-genome of different strain sets, we created multiple SNP sets by using different strain sets when perform analysis in various purposes. Totally 462,214 SNPs were identified from all 1,103 genomes, 650,683 SNPs were from 469 non-redundancy genomes, 355-8,921 SNPs were separately from 13 clonal groups.

### Population structure

The NJ trees were built by using the TreeBest software (http://treesoft.sourceforge.net/treebest.shtml) based on sequences of concatenated SNPs, and were visualized by using online tool iTOL [29].

The population structure of *V. parahaemolyticus* was built based on the 469 non-redundancy genomes set by using Chromosome painting and fineSTRUCTURE [17] as described before [16]. The fineSTRUCTURE result of all 469 non-redundancy genomes revealed that multiple clonal signals still presented. Therefore we selected only one representative genome from each clone, and combined them with the left genomes to perform another round of Chromosome painting and fineSTRUCTURE analysis. After six iterations, we finally obtained a set of 260 genomes with no clonal signals presented in the result (Supplementary Fig. 2b). To balance the sampling size among different populations, we selected 60 strains, including 14-16 strains from each population and 2 hybrid strains, to repeat the fineSTRUCTURE analysis (Supplementary Fig. 2c). The results further verify the population structure of *V. parahaemolyticus* species. Population assignment based on fineSTRUCTURE was consist with NJ tree (Fig. 1a) except for two strains, PCV08-7 and TUMSAT_H01_S4, and one epidemic group, CG2. Strain PCV08-7 and TUMSAT_H01_S4 were assigned to VppAsia and VppUS2 respectively by fineSTRUCTURE analysis, but in the NJ tree they are more closely related with VppX strains. The CG2 strains were all assigned to VppX populations by fineSTRUCTURE, but in NJ tree it was grouped with VppAsia strains. The length of chunks were extracted from the output file of Chromosome painting based on 469 non-redundancy strains, to calculate the percentage of admixtures of different populations for each non-redundancy genome (Fig. 2b).

### New designation of *V. parahaemolyticus* populations

In previous study, we designated four *V. parahaemolyticus* populations, named Asia-pop, US-pop 1, Hyb-pop 1, and Hyb-pop 2, separately, based on dataset of 157 genomes [16]. Here with more samples were used in distinguishing the population, we found a new population that mostly isolated from US, and the previously defined Hyb-pop 1 were known as just several hybrid strains, or representatives of otherwise unsampled populations. As evidences revealed that *V. parahaemolyticus* populations are geographical clustered, we proposed novel nomenclatures for them, which read as VppAsia, VppX, VppUS1 and VppUS2. The ‘Vpp’ is abbreviation of ‘*V. parahaemolyticus* population’. The first three populations are corresponding with previously defined Asia-pop, Hyb-pop 2 and US-pop 1, and the VppUS2 is the population newly identified in this study.

### Inference of substitution rate using BEAST

Two clonal groups with large sample size, CG1 (n = 153, global pandemic group, also known as O3:K6 and its sero-variants group) and CG2 (n = 92, an epidemic group that popular in US, also known as serotype O4:K12), were selected to calculate molecular clock respectively by using BEAST v1.83 [22]. The variations that caused by recombination inferred by our pipeline were excluded in substitution rates analysis. There are 10 of total 153 CG1 strains revealed too many strain-specific SNPs and revealed unusual long branches in the NJ tree even after removing the recombination variations (Supplementary Fig. 8), and similar pattern was observed in 1 of total 92 CG2 strains. These 11 strains with unusual high number of SNPs, and 22 strains with unknown isolation time, were excluded from BEAST analysis. We implemented analysis under GTR +Γ substitution model and relaxed clock model with constant size coalescent. The MCMC chain was run for 10^8^ and sampling for every 5,000 generations. The effective sample sizes of all inferred parameters were above than 200 in our results. The estimated molecular clock based on CG1 genomes is 5.6 × 10^-7^ with 95% confidence interval (CI) of 4.3-6.7 × 10^-7^ per site per year, and 5.4 × 10^-7^ with 95% CI of 3.6-7.2 × 10^-7^ for CG2 genomes. Here the average value, 5.5 × 10^-7^, was used as the most likely estimate of *V. parahaemolyticus* molecular clock. The extremes of 95% CI based on two clonal groups, i.e., 3.6-7.2 × 10^-7^, were used as lower and upper 95% boundaries for ensuring that they encompassed the true values as much as possible.

### Recombination detection and inference of recombination rate

Totally 13 clonal groups with more than 10 strains (Supplementary Table 1), defined by intra-group paired-distance less than 2,000 SNPs, were selected to be used in detection of recombination events. We firstly used previously pipeline to detect recombination [16]. Briefly, we recalled the SNPs for each clonal group because different datasets had different core-genomes, and these SNPs were used to construct a NJ tree. Then PAML software package [30] was used to determine the SNPs of each branch. Assuming neutrality and no recombination, the observed SNP density of a given region should follow the binomial distribution. We used the sliding window method to identify regions that rejected the null hypothesis (*P* < 0.05) and all SNPs in such windows were treated as recombined SNPs. We also used ClonalFrameML [31], a software based on maximum likelihood method, to detect bacterial recombination within the same dataset. Sequence alignments of genomes for each clonal group and the corresponding maximum-likelihood tree constructed using PHYML with HKY model [32], were used as input files and non-core regions were ignored during calculation. The inferred recombination regions using ClonalFrameML are mostly consistent with our in-house method (Supplementary Fig. 7).

Two sets of r/μ (ratio of size of recombination regions to the number of mutation sites) were obtained through different methods. The value is 331 with 99% CI of 228-435 for our in-house method and 295 with 99% CI of 186-404 for ClonalFrameML. The average value, 313, was selected as r/μ of the *V. parahaemolyticus*. Concerning the molecular clock rate of 5.5 × 10^-7^ per site per year, the most likely recombination rate of *V. parahaemolyticus* is 1.7 × 10^-4^ per site per year. We selected the extremes of r/ μ from two sets of 99% CI, 186-435, to calculate the lower and upper boundaries of recombination rate through multiplying the extremes of the molecular clock, which obtained the results of 6.7 × 10^-5^ - 3.1 × 10^-4^ per site per year. Accordingly, the time to obtained 1.01% of genome fragments would be 59 years with extreme boundaries of 32-151 years.

### Identify the contribution of *V. parahaemolyticus* populations to pandemic genomes in different geographical location

We assigned CG1 strains into two groups according to their isolated location, with one isolated from Asia and another isolated from North America. By using ClonalFrameML [31], we inferred the recombination fragments that occurred on each strain. Totally 81 fragments were found in Asia CG1 strains and 65 fragments were found in North America CG1 strains, with total size of 221 kb and the median length of 1,035 bp. Then we identify the possible donor genome of these recombination fragments by align them against with 468 non-redundancy genomes (excluding the CG1 genome from the dataset) using BLASTN, with a threshold of coverage >= 80% and identity >= 99.5%. The observed frequency of the donor genomes in each population was calculated. For recombination fragments carried by Asia CG1 genomes, the average value of their donor frequency in a population were taken as contribution proportion of the corresponding population to the recipient sequences in Asia CG1 genomes, and similarly, we obtained the contribution proportion of each population to the North America CG1 genomes (Fig. 3c). We also try the relaxed identity thresholds (99.0%) in BLASTN and acquired similar results.

## Acknowledgements

We gratefully acknowledge Dr. Narjol Gonzalez-Escalona for contributing *V. parahaemolyticus* genomes. This work is supported by the National Key Research & Development Program of China (No. 2017YFC1601503, 2016YFC1200100 and 2017YFC1200800), Sanming Project of Medicine in Shenzhen (No. SZSM201811071) and the National Natural Science Foundation of China (No. 31770001). J. Martinez-Urtaza were funded by Natural Environment Research Council (NERC) project (No. NE/P004121/1). D.F. is funded by a Medical Research Council Fellowship as part of the MRC CLIMB consortium for microbial bioinformatics (grant number MR/M501608/1).

## Author Contributions

Y. C., D. F. and R. Y. designed the study and coordinated the project; X. P., L. Y., J. M., Q. H., and D. Y. contributed strains for analysis; C. Y., X. P., Y. W, N. C., Y.Q. S., Y.J. S., Y. Y., M. J., Q., D. F. and Y. C. analyzed the data; E. F., J. M. and D. Z. provided insightful comments, F. and Y. C. wrote the manuscript. All authors approved the final version of the manuscript.

## References

1. Brown MV, Ostrowski M, Grzymski JJ, Lauro FM: A trait based perspective on the biogeography of common and abundant marine bacterioplankton clades. Mar Genomics 2014, 15:17–28.

2. Yilmaz P, Yarza P, Rapp JZ, Glockner FO: Expanding the World of Marine Bacterial and Archaeal Clades. Front Microbiol 2015, 6:1524.

3. Kent AG, Dupont CL, Yooseph S, Martiny AC: Global biogeography of Prochlorococcus genome diversity in the surface ocean. ISME J 2016, 10:1856–1865.

4. Hellweger FL, van Sebille E, Calfee BC, Chandler JW, Zinser ER, Swan BK, Fredrick ND: The Role of Ocean Currents in the Temperature Selection of Plankton: Insights from an Individual-Based Model. PLoS One 2016, 11:e0167010.

5. Nair GB, Ramamurthy T, Bhattacharya SK, Dutta B, Takeda Y, Sack DA: Global dissemination of Vibrio parahaemolyticus serotype O3:K6 and its serovariants. Clin Microbiol Rev 2007, 20:39–48.

6. Mutreja A, Kim DW, Thomson NR, Connor TR, Lee JH, Kariuki S, Croucher NJ, Choi SY, Harris SR, Lebens M, et al: Evidence for several waves of global transmission in the seventh cholera pandemic. Nature 2011, 477:462–465.

7. Weill FX, Domman D, Njamkepo E, Tarr C, Rauzier J, Fawal N, Keddy KH, Salje H, Moore S, Mukhopadhyay AK, et al: Genomic history of the seventh pandemic of cholera in Africa. Science 2017, 358:785–789.

8. Martinez-Urtaza J, van Aerle R, Abanto M, Haendiges J, Myers RA, Trinanes J, Baker- Austin C, Gonzalez-Escalona N: Genomic Variation and Evolution of Vibrio parahaemolyticus ST36 over the Course of a Transcontinental Epidemic Expansion. MBio 2017, 8.

9. Chin CS, Sorenson J, Harris JB, Robins WP, Charles RC, Jean-Charles RR, Bullard J, Webster DR, Kasarskis A, Peluso P, et al: The origin of the Haitian cholera outbreak strain. N Engl J Med 2011, 364:33–42.

10. Yeung PS, Boor KJ: Epidemiology, pathogenesis, and prevention of foodborne Vibrio parahaemolyticus infections. Foodborne Pathog Dis 2004, 1:74–88.

11. Su YC, Liu C: Vibrio parahaemolyticus: a concern of seafood safety. Food Microbiol 2007, 24:549–558.

12. Ansede-Bermejo J, Gavilan RG, Trinanes J, Espejo RT, Martinez-Urtaza J: Origins and colonization history of pandemic Vibrio parahaemolyticus in South America. Mol Ecol 2010, 19:3924–3937.

13. Martinez-Urtaza J, Trinanes J, Gonzalez-Escalona N, Baker-Austin C: Is El Nino a long-distance corridor for waterborne disease? Nat Microbiol 2016, 1:16018.

14. Baker-Austin C, Trinanes J, Gonzalez-Escalona N, Martinez-Urtaza J: Non-Cholera Vibrios: The Microbial Barometer of Climate Change. Trends Microbiol 2017, 25:76–84.

15. Yan Y, Cui Y, Han H, Xiao X, Wong HC, Tan Y, Guo Z, Liu X, Yang R, Zhou D: Extended MLST-based population genetics and phylogeny of Vibrio parahaemolyticus with high levels of recombination. Int J Food Microbiol 2011, 145:106–112.

16. Cui Y, Yang X, Didelot X, Guo C, Li D, Yan Y, Zhang Y, Yuan Y, Yang H, Wang J, et al: Epidemic Clones, Oceanic Gene Pools, and Eco-LD in the Free Living Marine Pathogen Vibrio parahaemolyticus. Mol Biol Evol 2015, 32:1396–1410.

17. Lawson DJ, Hellenthal G, Myers S, Falush D: Inference of population structure using dense haplotype data. PLoS Genet 2012, 8:e1002453.

18. Rosenberg NA, Pritchard JK, Weber JL, Cann HM, Kidd KK, Zhivotovsky LA, Feldman MW: Genetic structure of human populations. Science 2002, 298:2381–2385.

19. Wright S: Evolution in Mendelian Populations. Genetics 1931, 16:97–159.

20. Whitlock MC, McCauley DE: Indirect measures of gene flow and migration: FST not equal to 1/(4Nm + 1). Heredity (Edinb) 1999, 82 (Pt 2):117–125.

21. Didelot X, Meric G, Falush D, Darling AE: Impact of homologous and non-homologous recombination in the genomic evolution of Escherichia coli. BMC Genomics 2012, 13:256.

22. Drummond AJ, Suchard MA, Xie D, Rambaut A: Bayesian phylogenetics with BEAUti and the BEAST 1.7. Mol Biol Evol 2012, 29:1969–1973.

23. Ruiz GM, Rawlings TK, Dobbs FC, Drake LA, Mullady T, Huq A, Colwell RR: Global spread of microorganisms by ships. Nature 2000, 408:49–50.

24. Martinez-Urtaza J, Baker-Austin C, Jones JL, Newton AE, Gonzalez-Aviles GD, DePaola A: Spread of Pacific Northwest Vibrio parahaemolyticus strain. N Engl J Med 2013, 369:1573–1574.

25. Martinez-Urtaza J, Blanco-Abad V, Rodriguez-Castro A, Ansede-Bermejo J, Miranda A, Rodriguez-Alvarez MX: Ecological determinants of the occurrence and dynamics of Vibrio parahaemolyticus in offshore areas. ISME J 2012, 6:994–1006.

26. Chiesa S, Lucentini L, Freitas R, Nonnis Marzano F, Breda S, Figueira E, Caill-Milly N, Herbert RJH, Soares AMVM, Argese E: A history of invasion: COI phylogeny of Manila clam Ruditapes philippinarum in Europe. Fisheries Research 2017, 186:25–35.

27. Luo R, Liu B, Xie Y, Li Z, Huang W, Yuan J, He G, Chen Y, Pan Q, Liu Y, et al: SOAPdenovo2: an empirically improved memory-efficient short-read de novo assembler. Gigascience 2012, 1:18.

28. Delcher AL, Salzberg SL, Phillippy AM: Using MUMmer to identify similar regions in large sequence sets. Curr Protoc Bioinformatics 2003, Chapter 10:Unit 10 13.

29. Letunic I, Bork P: Interactive tree of life (iTOL) v3: an online tool for the display and annotation of phylogenetic and other trees. Nucleic Acids Res 2016, 44:W242–245.

30. Yang Z: PAML 4: phylogenetic analysis by maximum likelihood. Mol Biol Evol 2007, 24:1586–1591.

31. Didelot X, Wilson DJ: ClonalFrameML: efficient inference of recombination in whole bacterial genomes. PLoS Comput Biol 2015, 11:e1004041.

32. Guindon S, Gascuel O: A simple, fast, and accurate algorithm to estimate large phylogenies by maximum likelihood. Syst Biol 2003, 52:696–704.

