## Supplementary Figures for "Recent mixing of *Vibrio parahaemolyticus* populations"

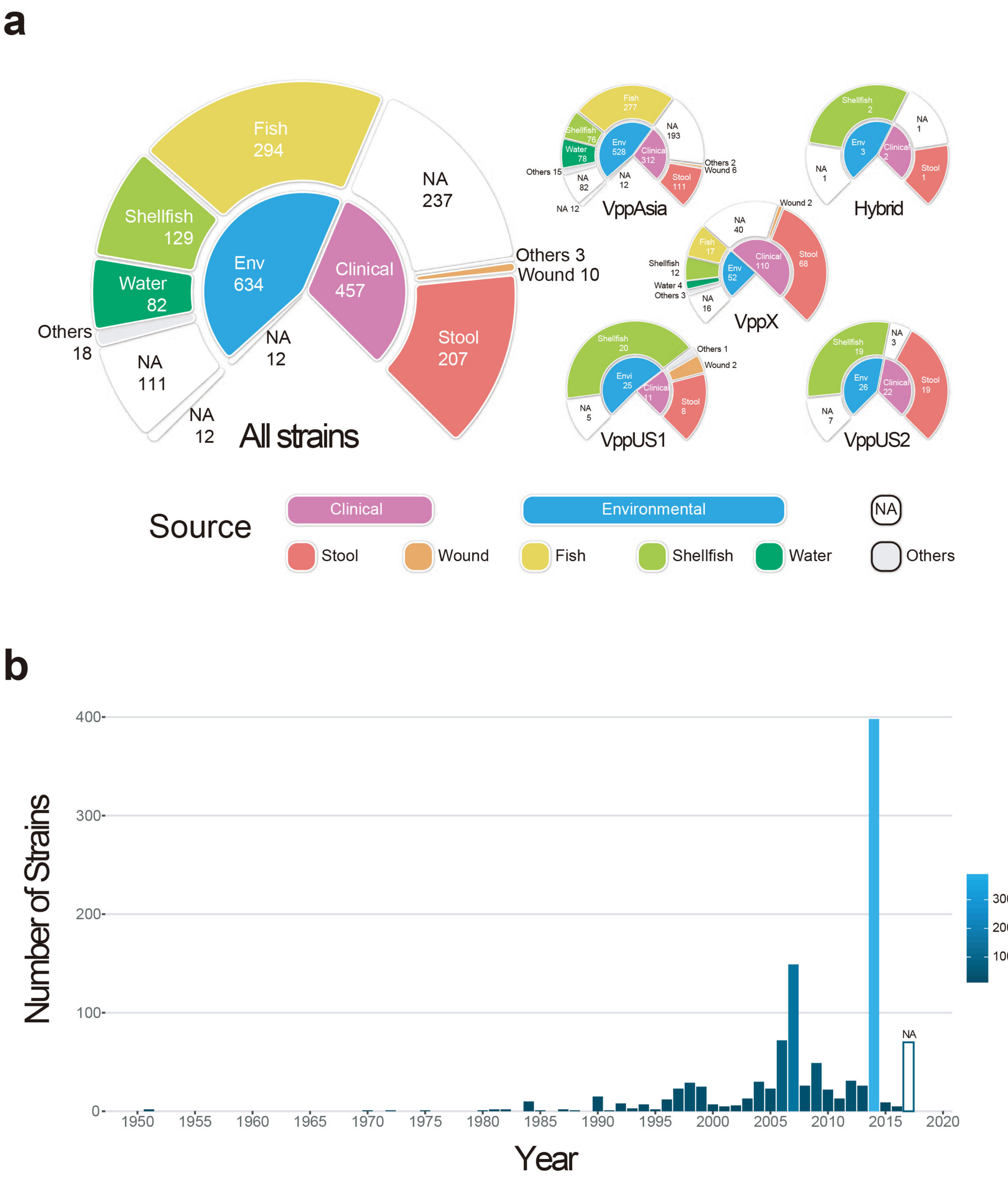


**Supplementary Figure 1.** Source (a) and isolation time (b) of 1,103 *V. parahaemolyticus* strains.

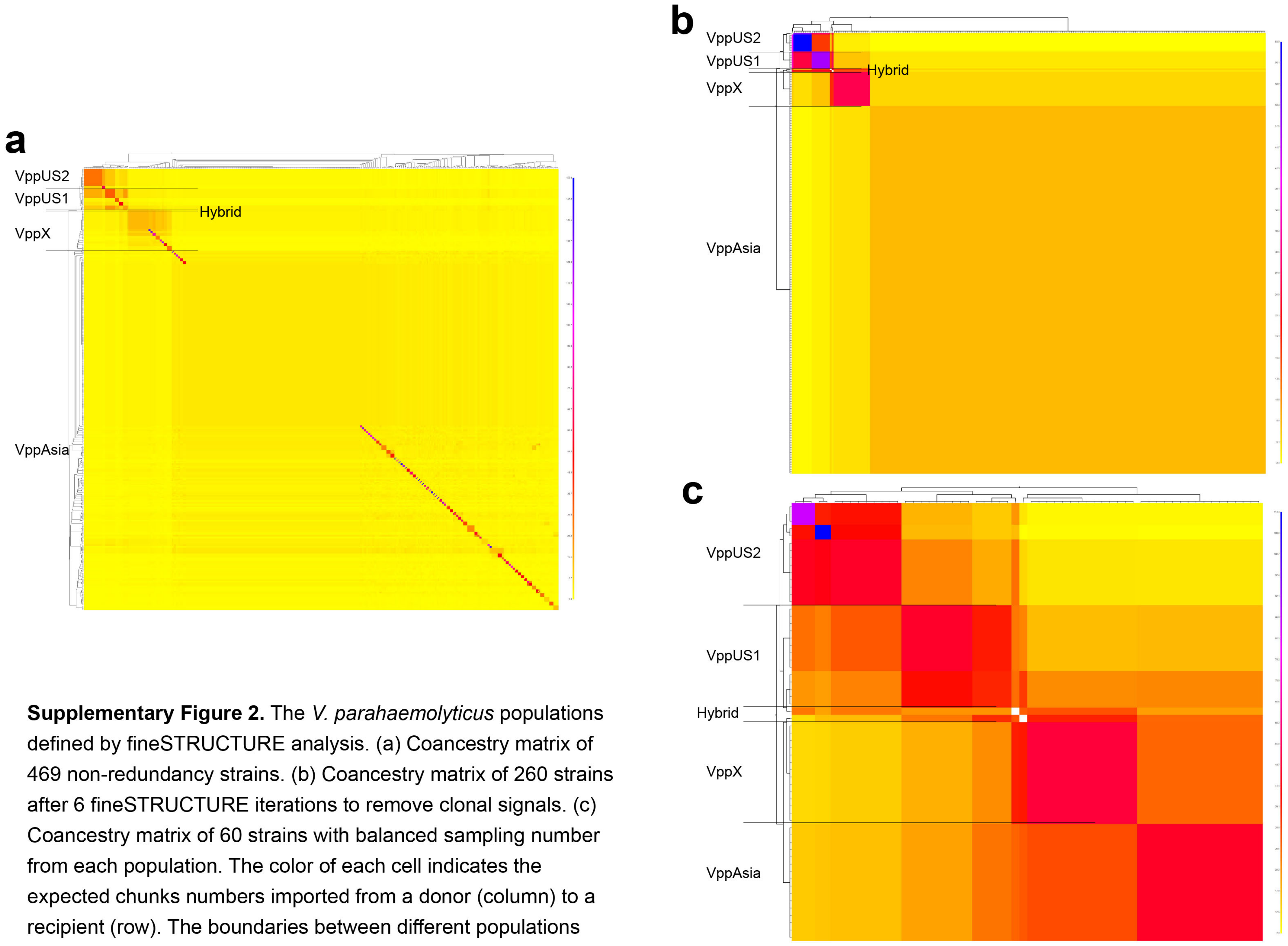


**Supplementary Figure 2.** The *V. parahaemolyticus* populations defined by fineSTRUCTURE analysis. (a) Coancestry matrix of 469 non-redundancy strains. (b) Coancestry matrix of 260 strains after 6 fineSTRUCTURE iterations to remove clonal signals. (c) Coancestry matrix of 60 strains with balanced sampling number from each population. The color of each cell indicates the expected chunks numbers imported from a donor (column) to a recipient (row). The boundaries between different populations are marked with lines.

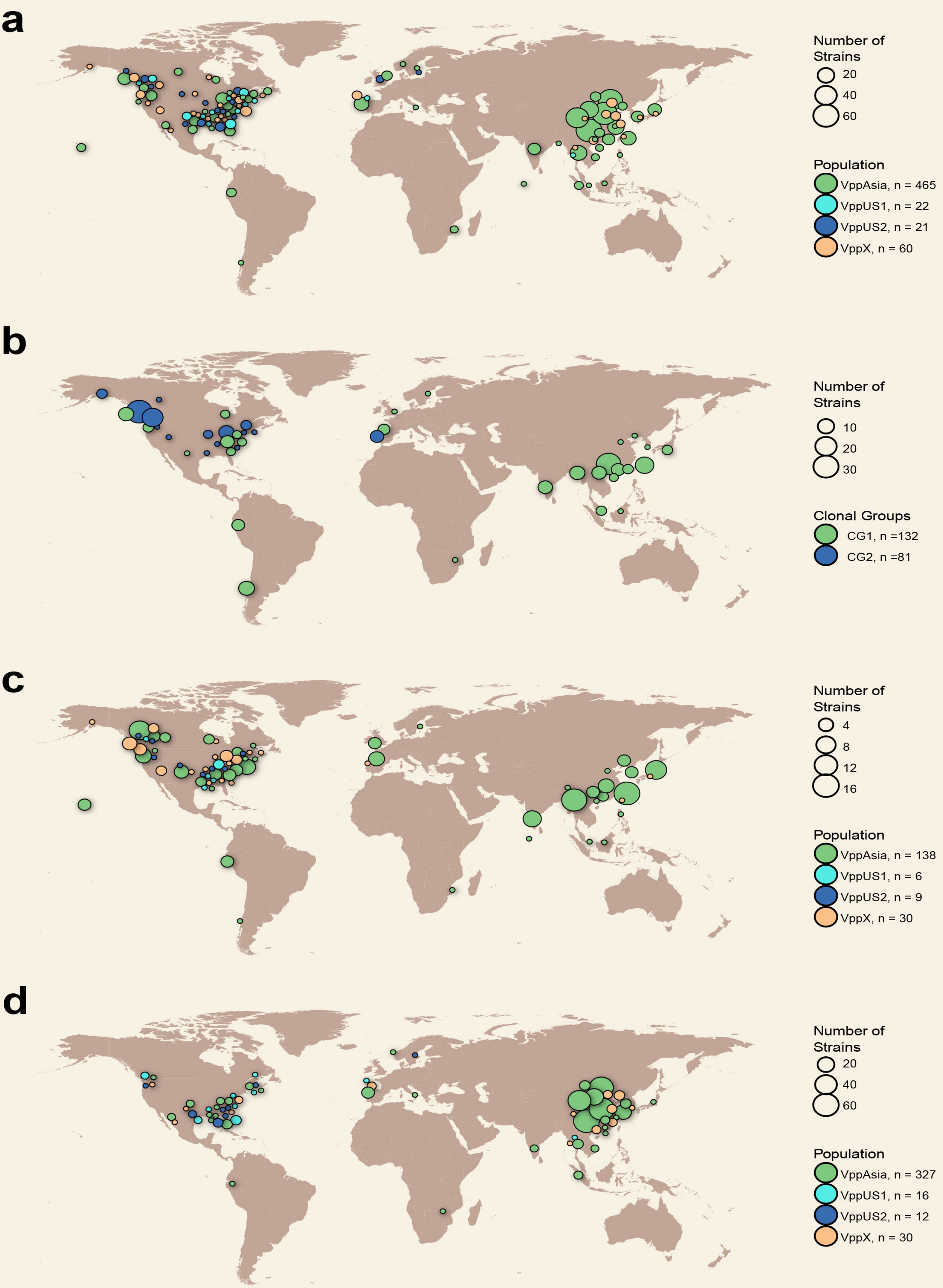


**Supplementary Figure 3.** Geographical distribution of *V. parahaemolyticus* populations (a), two major clonal groups (b), clinical samples (c) and environmental samples (d). Some samples that isolated from a same city (or country if there are no city information) belong to a same clonal group, which will influence the visual effect of the population distribution. Therefore, the clone within same city/country were counted as one strain in panel a, c and d.

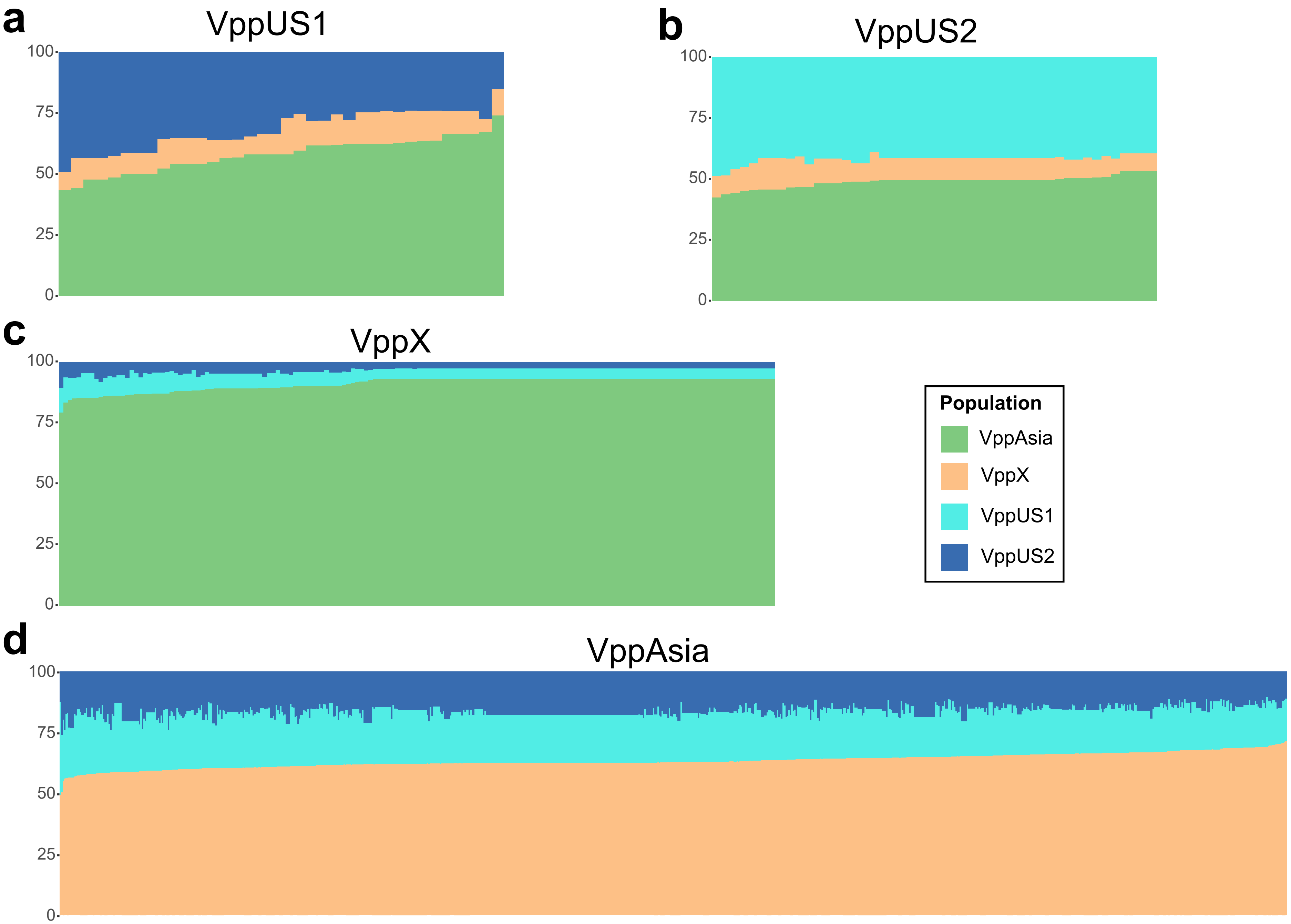


**Supplementary Figure 4.** The genome admixture of four *V. parahaemolyticus* populations according to chromosome painting. Chromosome painting results for all VppUS1 (a), VppUS2 (b), VppX (c) and VppAsia (d) strains inferred by using other populations as donor. Each vertical bar represents one isolate and the proportion indicates the contribution of each population.

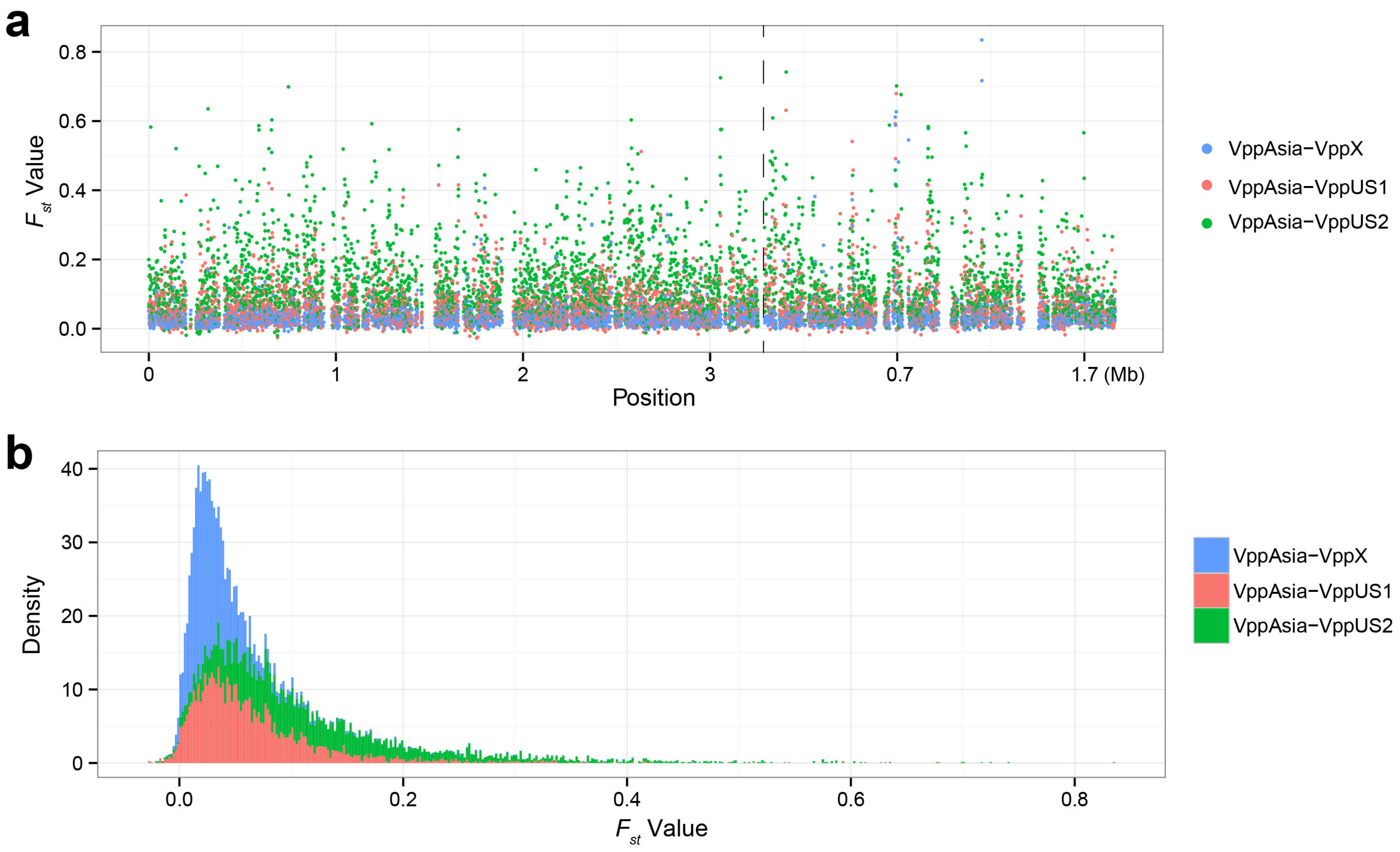


**Supplementary Figure 5.** $F_{st}$ value sliding window analysis (a) and overall distribution (b) between VppAsia and other populations. Window size and step size were set to 1 Kb. Two chromosomes are separated by dot line in panel a.

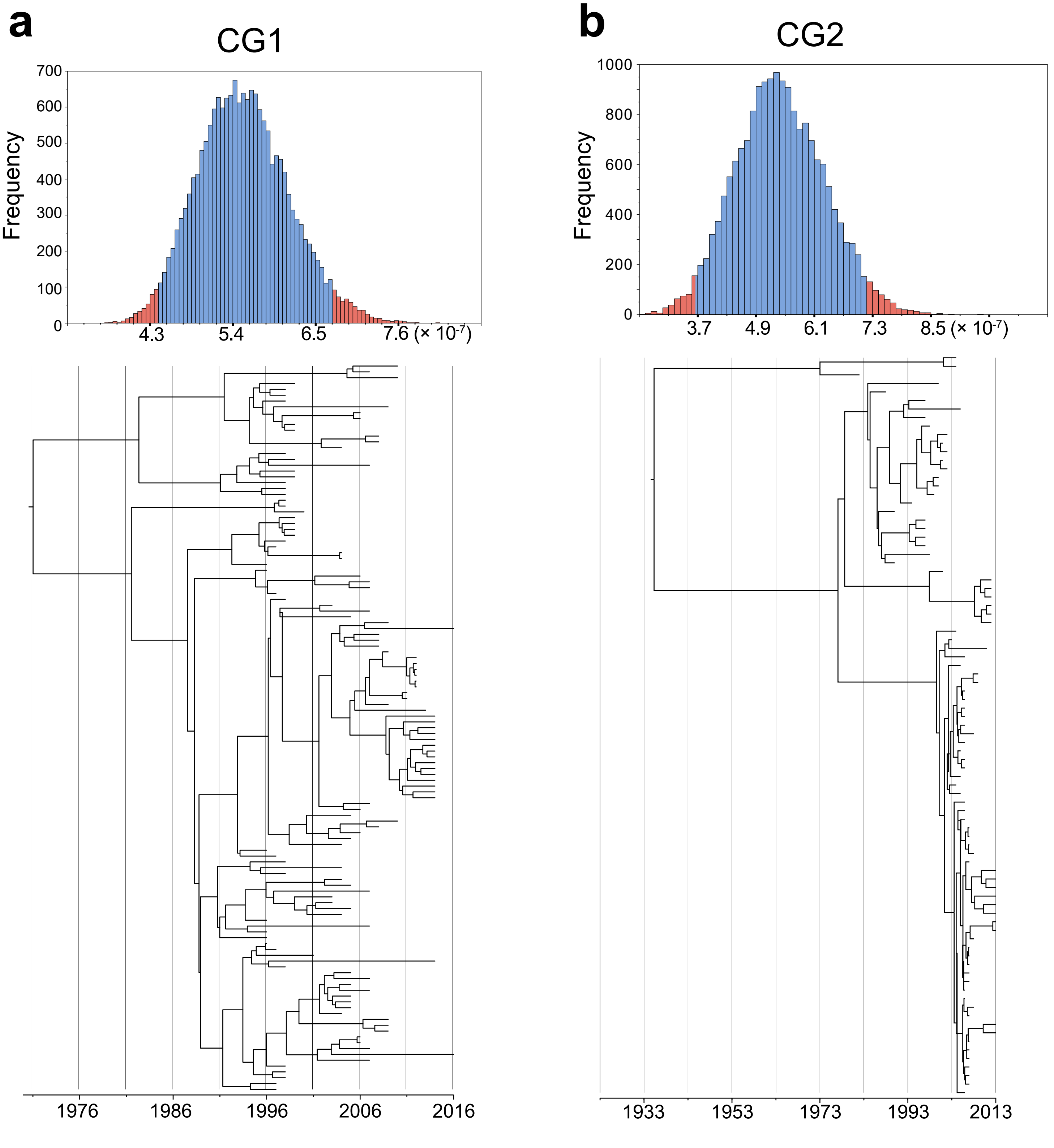


**Supplementary Figure 6.** Clock rate of two major clonal groups CG1 (a) and CG2 (b) inferred by BEAST. Top: the posterior probability density distribution of clock rate. Bottom: maximum clade credibility tree.

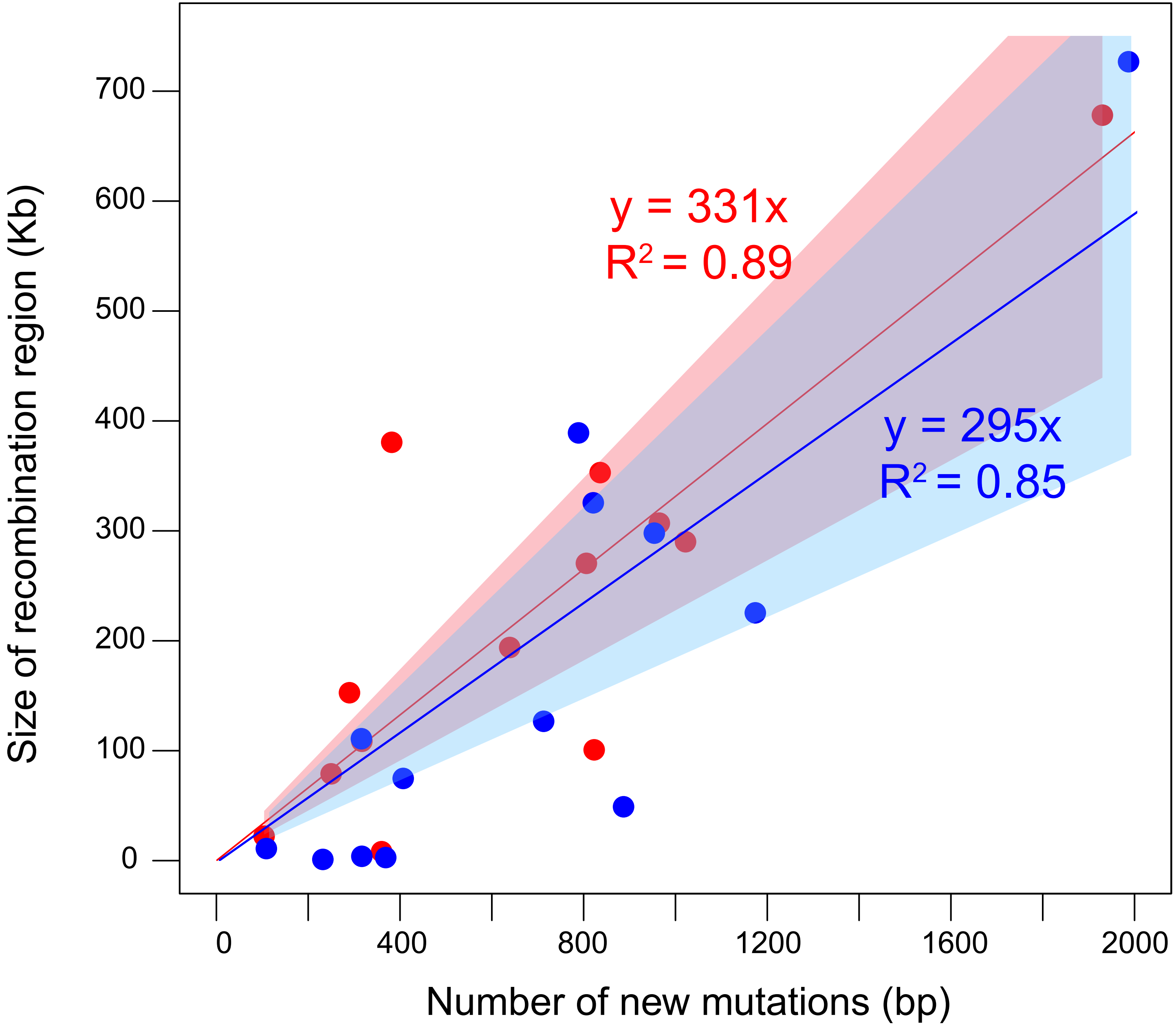


**Supplementary Figure 7.** Correlation between total size of recombination regions and amount of new mutation sites within 13 clonal groups. The red color indicates results that inferred by pipeline in previous paper, blue indicates results inferred by ClonalFrameML. Each dot indicated one clonal lineage. The linear regression was constrained to go through the origin and the light red and blue shading indicates the 99% CI of the slope, [228-435] for red and [186-404] for blue.

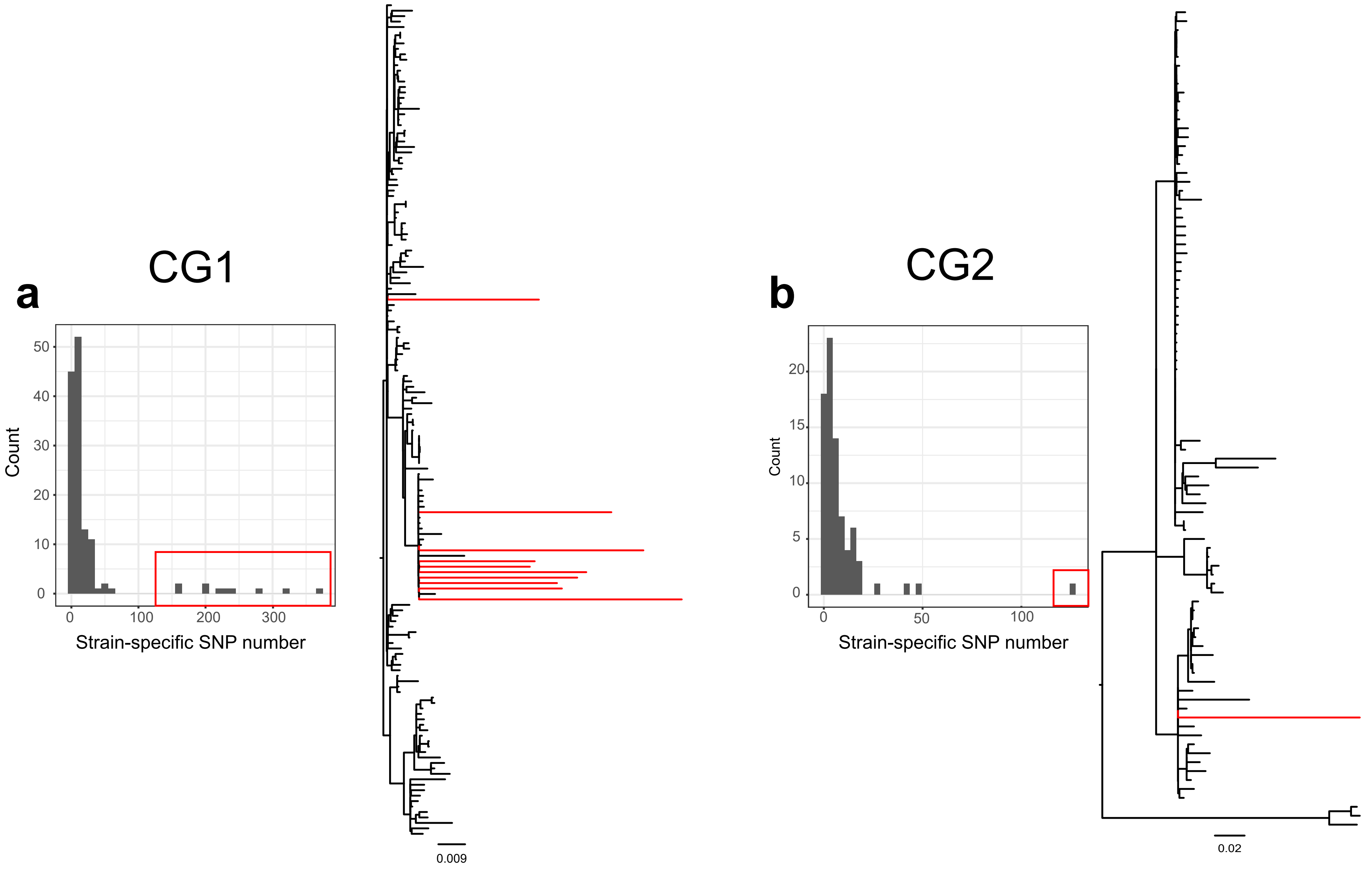


**Supplementary Figure 8.** *V. parahaemolyticus* strains contained unusual high number of strain-specific variations. The NJ trees of CG1 (a) and CG2 (b) were build based on SNPs after removing recombination sites. Both the long branches in NJ tree and outliers in the histograms indicated some *V. parahaemolyticus* strains contained unusual high number of strain-specific SNPs. Strains marked in red in histogram/NJ tree are excluded from BEAST analysis.

**Supplementary Table 1.** Background information of 1,103 strains used in this research (seperate file).

**Supplementary Table 2.** The nucleotide differences (green) and $F_{st}$ value (yellow) among *V. parahaemolyticus* populations.

| | VppAsia | VppX | VppUS1 | VppUS2 |
|---|---|---|---|---|
| VppAsia | 30227 | 31168 | 33325 | 34357 |
| VppX | 0.039 | 29729 | 33515 | 34587 |
| VppUS1 | 0.073 | 0.085 | 31637 | 33004 |
| VppUS2 | 0.138 | 0.151 | 0.082 | 28964 |
